# Optogenetic Regulation of Mitochondrial Function to Modulate Cell Death

**DOI:** 10.64898/2026.07.19.739443

**Authors:** Run-Zhou Yang, Dian-Dian Wang, Sen-Miao Li, Dan-Hua Liu, Pei-Pei Liu, Shu-Ang Li, Jian-Sheng Kang

## Abstract

Cell death is a critical process involved in physiological and pathological conditions, including neurodegenerative diseases and cancer. This study explores the use of optogenetic techniques to induce cell death by employing light sensitive proteins. By manipulating mitochondrial function with light-sensitive proteins, we investigated three distinct strategies: 1) inhibiting oxidative phosphorylation through Gloeobacter rhodopsin-mediated alkalization, 2) inducing mitochondrial depolarization with reverse proton-pumping rhodopsins (RPPR) and anion-conducting channelrhodopsins, and 3) generating reactive oxygen species (ROS) using mitochondria-targeted miniSOG. Our findings highlight the potential of optogenetic approaches to induce cell death, offering promising avenues for therapeutic interventions in diseases characterized by aberrant cell survival.

## Introduction

Cell death is the irreversible loss of cellular function, which can be caused by external factors like physical or chemical stimuli, hypoxia, trauma, viral infection, or spontaneous processes such as embryonic development and nervous system maturation^1^. Cell death modulation is crucial for physiological and pathological processes. Excessive cell death is associated with neurodegenerative diseases like Alzheimer’s and Parkinson’s^2,3^, while cancer cells evade apoptosis to maintain uncontrolled proliferation ^4^. Cell death can be triggered by various physiological factors, including abnormal pH^5^, deceased mitochondrial membrane potential^6^ and excessive reactive oxygen species^7^. The regulation of intracellular pH is essential for many cellular processes, including enzymatic reactions and signal transduction ^8^. Acidification of the cytoplasm during ischemia can lead to cell swelling and necrosis ^9^. Conversely, alkalization of the cytoplasm could trigger the opening of mitochondrial permeability transition pore (MPTP)^10^, which exhibits pH-dependent opening above pH 7.3-7.5 and closure below pH 7, ultimately resulting in calcium overload and subsequent cell death^11,12^. Proton pumping rhodopsins (PPR), such as bacteriorhodopsin (BR), archaerhodopsin (Arch), and Gloeobacter rhodopsin (GR), actively transport protons from the intracellular environment to the extracellular space, resulting in a significant increase in cytosolic pH^13^. In contrast, reverse proton-pumping rhodopsins (RPPR) transport protons from the extracellular environment into the cell. The first RPPR was engineered from a non-ion conducting sensory rhodopsin, ASR, through a single site mutation, D217E^14^. Besides ASR(D217E), reverse proton pumps also exist in nature, including PoXeR^15^, NsXeR^16^, and AntR^17^.

Mitochondrial membrane potential (Δψ_m_) depolarization is associated with apoptosis induction^18^ and selective removal of damaged mitochondria through mitophagy^19^. Natural chloride ion channels have been identified in mitochondria, where they play a role in apoptosis induction^20^. Anion-conducting channelrhodopsins, such as iChloC^21^, iC1C2^22^, GtACR1^23^ and GtACR2^24^ are cation-conducting light-gated channels. Our previous work discovered that GtACR1 can be targeted to the mitochondria in astrocytes and induced dopaminergic neuronal death^25^. The targeting of GtACR1 to mitochondria for inducing cell death has not yet been reported.

Reactive oxygen species (ROS), which generated as byproducts of oxidative phosphorylation during the electron transport chain^26^, could act as signaling molecules under normal conditions^27^ but cause cell damage and apoptosis when accumulated excessively^28^. ROS generation can be resulted from both mitochondrial depolarization and hyperpolarization^29,30^. Chromophore molecules that generate ROS can also be used to induce cell death^31,32^. MiniSOG, a small flavoprotein from *Arabidopsis thaliana*, generates singlet oxygen when illuminated by blue light ^33^.

In this paper, we discuss various strategies of manipulating mitochondrial function for inducing cell death utilizing light-sensitive proteins. These include the inhibition of oxidative phosphorylation through GR-induced alkalization, mitochondrial depolarization through mt-ASR(D217E) and mt-ACR1, and generation of free radicals via mt-miniSOG.

## Results

Since intracellular pH is tightly linked to energy metabolism and cell fate, we asked whether an optogenetically imposed alkalinization would be sufficient to compromise ATP production and trigger cell death. To explore the influence of light induced intracellular pH changes on cell survival, a proton pumping rhodopsin derived from Gloeobacter (GR) was expressed on the plasma membrane of HeLa cells (Figure 1). Cells transfected with pHluorin-tagged GR exhibited an increase in intracellular pH upon light stimulation, as evidenced by the fluorescence increasing of pHluorin after light exposure (Figure 1A, Figure S1A). In contrast, cells transfected with pHluorin-tagged non-pumping mutant GR(D121N) demonstrated no obvious change of fluorescence. These results were consistent with those obtained using SNARF-1-AM, a cell-permeable ratiometric pH-sensitive dye (Figure 1A, second row and Figure 1B, Supplementary Movie 1). Whole-cell patch clamping recordings revealed that GR generated an outward current upon green laser stimulation (Figure 1C). Upon light exposure, *E. coli* expressing GR significantly lowered the pH of unbuffered suspensions, confirming GR’s role as a light-activated proton pump (Figure S1B). Intriguingly, by fusing the C terminal of GR with a ATP sensor, AT1.03^34^, a decrease in intracellular ATP upon light stimulation was observed (Figure 1E, dark vs. light: *p* < 0.0001, *t-test*, Supplementary Movie 2). In contrast, cells expressing the non-pumping mutant GR(D121N) exhibited no ATP alterations (Figure 1D, dark vs. light: *p* > 0.05, *t-test*). Consistent with these observations, chemiluminescence-based ATP measurement showed reduced ATP levels in GR-expressing cells compared to GR(D121N)-expressing cells after light exposure (Figure 1F; GR: light vs. dark, *p* < 0.0001; GR(D121N), light vs. dark, *p* > 0.05). The ATP decreased is contributed to the alkalization effect of light-driven proton pumping, as another PPR, bacteriorhodopsin (BR) could also cause the decrease of intracellular ATP accompanied by an outward current and intracellular pH increasement upon light stimulation (Figure S2). Since ATP in mammalian cells is generated through glycolysis and oxidative phosphorylation, the pH-induced decrease in ATP could potentially inhibit one or both metabolic pathways. By blocking energy supply using glycolysis inhibitor 2DG and the oxidative phosphorylation inhibitor oligomycin, the light-induced ATP change was examined. When both inhibitors were used in combination, the pH-induced ATP decrease was blocked, whereas ATP levels declined under the individual application of either 2DG or oligomycin (Figure 1G; ATP decrease in each group, Δ mean ratio ± standard deviation (SD): control, -0.1114 ± 0.029; 2DG, -0.1527 ± 0.054; oligomycin, -0.2938 ± 0.072; 2DG + oligomycin, 0.0019 ± 0.031; control and 2DG + oligomycin, *t-test*, *p* = 0.0004; 2DG and 2DG + oligomycin, *t-test*, *p* = 0.0003; oligomycin and 2DG + oligomycin, *t-test*, *p* < 0.0001). This suggested that light exposure can suppress ATP production via glycolysis and oxidative phosphorylation pathways. Moreover, the oxygen consumption rate (OCR) decreased under light illumination in GR stably transfected HEK293t cells (Figure 1H). In contrast, control cells demonstrated no significant alterations in OCR upon light illumination (Figure 1H). This suggested that light can inhibit oxidative phosphorylation in GR stably transfected HEK293t cells. As ATP is crucial for cell survival, light-induced ATP decrease by GR could potentially induce cell death. As a test of this proposal, GR was expressed on the plasma membrane of primary cultured cardiomyocytes. Propidium Iodide (PI) is a fluorescent dye that assesses cell viability by penetrating compromised membranes of dead or late-stage apoptotic cells, binding to nucleic acids, and fluorescing red. Prolonged light stimulation caused morphological changes of the cell (Figure 1I, second row). The PI staining results demonstrated that light could trigger cell death in GR expressing cardiomyocytes (Figure 1I and 1J; n = 9 cells, *t-test*, *p* < 0.0001, Supplementary Movie 3). Moreover, cardiomyocytes without GR expression subjected to the same light stimulation protocol demonstrated no cell death, ruling out the possibility of light toxicity (n = 5 cells, *t-test*, *p* > 0.05).

**Figure 1.**
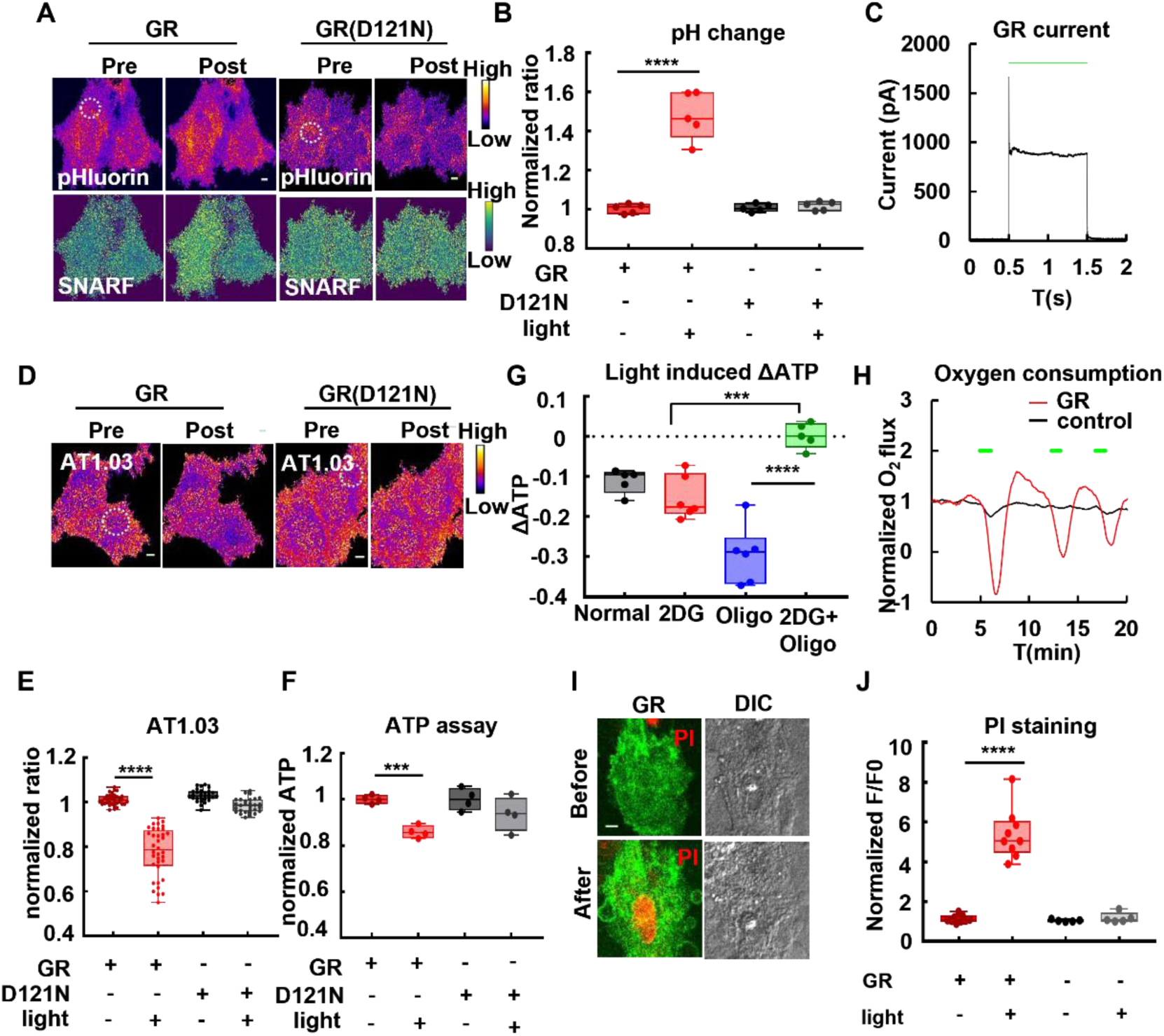
Light exposure induces alkalization, ATP decrease, and cell death in cells expressing membrane-targeted GR. **(A)** Light induces intracellular alkalization in GR expressing cells. HeLa cells transfected with GR-pHluorin or GR(D217N)-pHluorin were stained by SNARF-1 AM dye and subjected to light stimulation. The upper row indicated the pHluorin fluorescence and the lower row indicated the SNARF ratio. The images were represented as pseudo-color. The dashed white circle indicates the area with light stimulation (515 nm, 20 seconds). Scale bar, 5 μm. **(B)** Quantification of the SNARF ratio in (A). There was a significant difference between illuminated and non-illuminated GR expressing cells (n = 5 cells, *t-test*, *p* < 0.0001). **(C)** Patch clamping recording of photocurrent of HeLa cells expressing GR. **(D)** Light induced ATP decrease in cells expressing GR. HeLa cells expressing the fusion protein GR-AT01.03 or GR(D217N)-AT01.03 were subjected to light stimulation. The dashed white circle indicates the area with light stimulation (559 nm, 1.2 min). The intracellular ATP level was indicated by the ratio of AT1.03. The images were represented as pseudo-color. Scale bar, 5 μm. **(E)** Quantification of the AT1.03 ratio in (D). There was a significant decrease of AT1.03 ratio in GR-expressing cells, while GR(D121N)-expressing cells showed no significant change upon light stimulation (*t-test*, GR, n = 36 cells, *p* < 0.0001; GR(D217N), n = 29 cells, *p* > 0.05). **(F)** ATP assay measured by chemiluminescence in cells expressing GR and GR(D121N) before and after light exposure. Consistent with AT1.03 result, there was a significant decrease in ATP level in GR but not in GR(D121N) expressing cells (*t-test*, GR, n = 4 experiments, *p* < 0.0001; GR(D217N), n = 4 experiments, *p* > 0.05). **(G)** Amplitude of light-induced ATP decreases in GR-expressing cells pre-treated with inhibitors. In GR-expressing HeLa cells pretreated with oligomycin, the extent of light-induced ATP decrease was more pronounced than that under normal conditions (*t-test*, *p* < 0.005). In GR-expressing cells pre-treated with 2DG and oligomycin, light stimulation no longer caused a decrease in ATP (Normal, n = 5 cells; 2DG, n = 6 cells; Oligomycin, n = 6 cells; 2DG + Oligomycin, n = 5 cells). **(H)** Oxygen consumption rate (OCR) in GR-expressing and control cells upon light stimulation. The OCR curves of GR-expressing and control cells were represented as red and black, respectively. The green bars indicate light stimulation. **(I-J)** Light induces cell death in primary cardiomyocytes expressing GR. **(I)** Images of GR-expressing cardiomyocytes stained with PI before and after light stimulation. Scale bar, 5 μm. **(J)** Quantification of PI fluorescence in (I). There was a significant difference in cell death between illuminated and non-illuminated cells (n = 9 cells, *t-test*, *p* < 0.0001). In control cells exposed to light, there was no significant difference compared to cells kept in the dark (n = 5 cells, *t-test*, *p* > 0.05).

Mitochondrial membrane potential (MMP) plays a critical role in the regulation of cell death pathways. The loss of MMP can be a signal of bioenergetic stress, potentially resulting in the release of apoptotic factors that trigger cell death^35^. Uncoupling proteins (UCPs), located in the inner mitochondrial membrane, modulate the mitochondrial membrane potential (MMP) by allowing protons to bypass ATP synthase and thereby uncouple electron transport from ATP production, resulting in elevated heat generation^36^. Reverse proton-pumping rhodopsins (RPPR) pump proton from extracellular space into cell. By transferring RPPR to mitochondria, RPPR could function like uncoupling proteins. To target RPPR to mitochondria, a signal sequence derived from the 1-135 aa of ABCB10^37^, or a four-time repeated signal sequence of cytochrome c oxidase (4cox8) were fused with the N-terminal of three RPPRs, including ASR(D217E), PoXeR and NsXeR. The C-terminal of these RPPRs were fused to EGFP for monitoring their localizations. The efficiency of mitochondrial targeting was assessed by calculating the Pearson’s correlation coefficient between the mitochondria-targeted RPPRs and that of a mitochondrial marker, TMRM (Figure 2A). The results are as follows: ASR(D217E) showed a high correlation coefficient of 0.93 ± 0.03, indicating strong mitochondrial localization. PoXeR had a moderate correlation coefficient of 0.75 ± 0.17, suggesting partial mitochondrial targeting (Figure 2B, second row). In contrast, NsXeR exhibited a low correlation coefficient of 0.24 ± 0.52, which suggests a weak mitochondrial localization (Figure S3, first row). It is noteworthy that the targeting efficiency of the mt-fused NsXeR and PoXeR revealed two distinct patterns: one group with strong mitochondrial localization and another with non-mitochondrial localization. These two groups can be distinguished from the violin plot in Figure 2A. As depicted in Figure 2B and Figure S4, ASR(D217E) demonstrated successful mitochondrial localization in different types of cells, including HeLa, COS7 and HepG2. Therefore, ASR(D217E) was selected for further evaluations.

**Figure 2.**
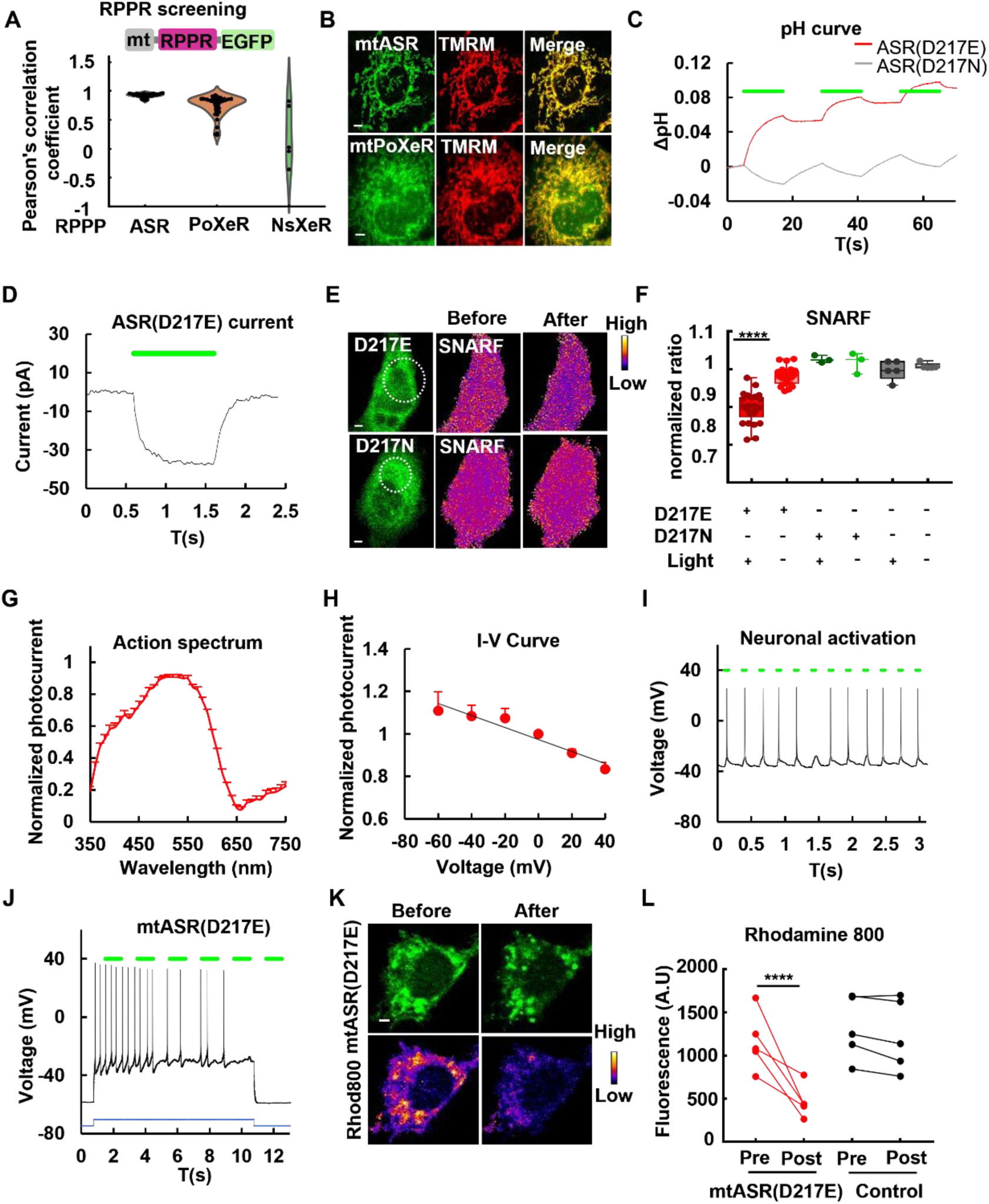
Screening and characterization mitochondrial-targeted reverse proton pumps. **(A)** Screening of mitochondrial-targetable reverse proton pumps. The N-termini of three reverse proton pumping rhodopsin (RPPR), including ASR(D217E), PoXeR and NsXeR were fused to mitochondrial targeting sequences (MTS), while their C-termini were fused to EGFP to enable visualization of their cellular localization. The mitochondrial localization of reverse proton pumps was assessed by Pearson correlation coefficients between fluorescent proteins and mitochondrial markers. **(B)** Confocal images of MTS fused ASR(D217E) and PoXeR expressed in HeLa cells. TMRM was used as a mitochondrial marker. Scale bar, 5 μm. **(C)** Light-induced pH change of *E.coli* suspension expressing ASR(D217E) or ASR(D217N). The curves of ASR(D217E) and ASR(D217N) are shown in red and gray, respectively. Green bars indicate light stimulation. **(D)** Patch-clamp recording of photocurrents in HEK293t cells expressing ASR(D217E). Green bar indicates light stimulation. **(E)** Confocal images of HeLa cells expressing ASR(D217E) or ASR(D217N) stained with SNARF before and after light stimulation. The white dotted circle indicates light stimulation area. The images were represented as pseudo-color. Scale bar, 5 μm. **(F)** Quantification of SNARF ratios in (E). ASR(D217E)-expressing cells exhibited significant decrease in SNARF ratios after light stimulation (n = 26 cells, paired *t-test*, *p* < 0.0001), while ASR(D217N)-expressing cells and control cells did not exhibit significant changes in SNARF ratios after light illumination (ASR(D217N), n = 3 cells, control, n = 5, paired *t-test*, *p* > 0.05). **(G)** Photocurrent spectrum of HEK293t cells expressing ASR(D217E). The peak of photocurrent was reached at 510 nm (n = 20 cells). **(H)** Normalized photocurrent of HEK293t cells expressing ASR(D217E) at different holding voltages (n = 7 cells). **(I)** Light induced action potential firing in primary neurons expressing ASR(D217E). The green bars indicate light stimulation. **(J)** Mitochondrial-targeted ASR(D217E) exhibits no plasma membrane leaky targeting. Light stimulation could not trigger action potential firing of neurons expressing mt-ASR(D217E). The blue curve indicates the injected current, and the green bars indicate light stimulation. **(K)** Confocal images of HeLa cells expressing mt-ASR(D217E) stained with rhodamine 800 before and after light stimulation. The rhodamine 800 fluorescence was represented as pseudo-color. Scale bar, 5 μm. **(L)** Quantification of rhodamine 800 fluorescence before and after light stimulation in mt-ASR(D217E)-expressing cells or control cells. There was a significant decrease of rhodamine 800 fluorescence in mt-ASR(D217E)-expressing cells upon light stimulation. In contrast, no significant difference in rhodamine 800 fluorescence was observed in control cells before and after light illumination (mt-ASR(D217E), paired *t-test*, n = 5 cells, *p* < 0.0001; control, paired *t-test*, n = 5 cells, *p* > 0.05).

The reverse proton pumping activity of ASR(D217E) was confirmed by expressing it in *E. coli*. Illumination of suspension of ASR(D217E)-expressing *E. coli* in non-buffering solution led to an increase in extracellular pH (Figure 2C, red curve). In contrast, no increase was observed in *E. coli* expressing ASR(D217N), a non-pumping mutant (Figure 2C, grey curve). A light induced inward current was recorded in ASR(D217E)-expressed HEK293t cells using patch clamping (Figure 2D). This inward current could also be observed in PoXeR-expressed HEK293t cells (Figure S5). Utilizing the pH sensitive dye, SNARF-1-AM, a decrease in intracellular pH was observed upon light stimulation in ASR(D217E)-expressing HeLa cells, but not in ASR(D121N)-expressing cells (Figure 2E-F; ASR(D217E): light vs. dark, *p* < 0.0001, *t-test*). Further characterization of ASR(D217E) in HEK293t revealed its action spectrum and I-V curve (Figure 2G-H). The maximum photocurrent was generated under 510 nm, and the photocurrent was found to be voltage sensitive, with enhanced response at lower voltage. The photocurrent of ASR(D217E) demonstrated a light intensity dependent manner, with a saturation photocurrent reached at more than 100 mW/cm^2^ (Figure S6A). Furthermore, ASR(D217E) could be used as an optogenetic activator to evoke action potentials through light stimulation in neurons (Figure 2I, Figure S6B). In contrast to ASR(D217E), mitochondrial targeted mt-ASR(D217E) failed to trigger action potentials, indicating no leaky expression on the plasma membrane of mt-ASR(D217E)-expressed cells (Figure 2J). Given that ASR(D217E) is voltage sensitive, the depolarization effect of ASR(D217E) on the mitochondrial membrane potential (-180 mV) should be greater than on the plasma membrane. This effect makes mt-ASR(D217E) a potential light activated MMP dissipation inducer. This was confirmed by a significant decrease in mitochondrial membrane potential, as measured with rhodamine 800, an MMP sensitive dye, upon light stimulation in mt-ASR(D217E) expressed HeLa cells (Figure 2K-L; mt-ASR(D217E): light vs. dark, *p* < 0.0001, *paired t-test*). In contrast, control cells demonstrated no change in MMP upon light stimulation (Figure 2L; control: light vs dark, *p* = 0.0672, *paired t-test*). Moreover, the morphology of mitochondria was altered after light stimulation, with more fragmented and granule-like structures observed (Figure 2K, second row).

Mitochondrial chloride conducting channels like CLC4^20^ and CLC5^38^ involved in the induction of cell death. To mimic the effects of these mitochondrial chloride conducting channels, light-gated anion-conducting channelrhodopsins were engineered to target to mitochondria, with a four-time repeated signal sequence of cytochrome c oxidase (4cox8) fused to the N-terminus of light-gated chloride channels. Their C-terminus were fused to EGFP for monitoring of targeting. Three light-gated chloride channels (ChloC), including iC1C2, ACR1, and ACR2 were investigated. The efficiency of mitochondrial targeting was assessed by calculating the Pearson’s correlation coefficient between the fluorescence of mitochondria-targeted light-gated chloride channels and that of a mitochondrial marker, TMRM (Figure 3A). The Pearson’s correlation coefficients are as follows: ACR1, 0.87 ± 0.079; ACR2, 0.50 ± 0.16; iC1C2, 0.16 ± 0.22. As shown in Figure 3A-B and Figure S7A, ACR1 exhibited successful mitochondrial localization, while ACR2 and iC1C2 failed to target to mitochondria. The localization of mt-ACR1 was further confirmed in various cell types, including HeLa cells, cardiomyocytes, and neurons (Figure 3B). To determine the optimal wavelength for light stimulation, action spectra were recorded by patch clamping in HEK293t cells. The peak of ACR1 photocurrent was achieved at around 510 nm, while ACR2 and iC1C2 were blue shifted, with peaks at 370 nm and 450 nm, respectively (Figure 3C). To explore the voltage sensitivity of light-gated chloride channels, I-V curves were recorded in HEK293t cells. The relationships between photocurrent and voltage appeared to be exponential in ACR1 and ACR2, and linear in iC1C2 (Figure S7B).

**Figure 3.**
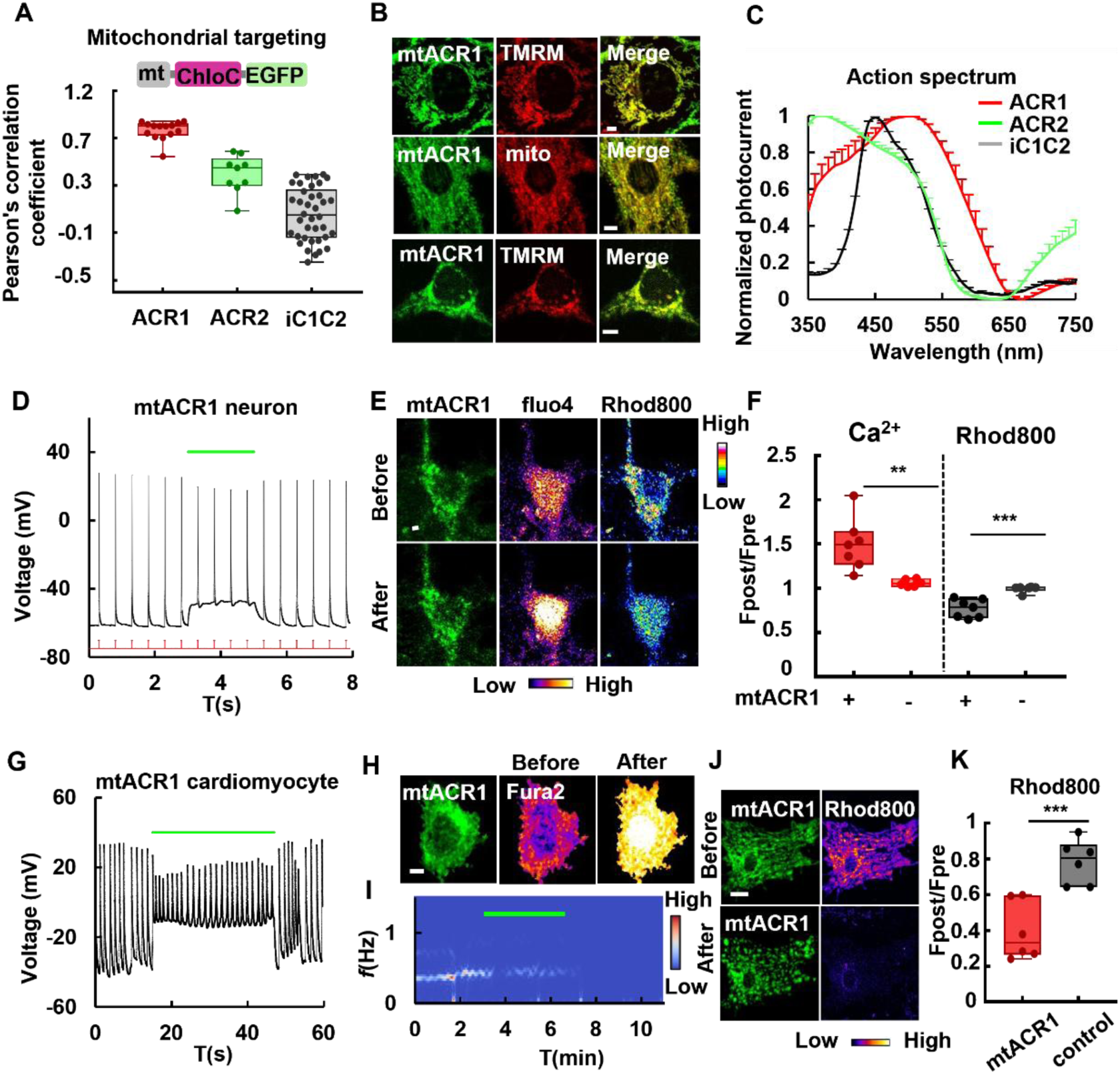
Characterization of mitochondrial-targeted chloride ion channels. **(A)** Screening of mitochondrial-targetable chloride ion channels. The N-termini of three chloride channels (ChloC), including ACR1, ACR2 and iC1C2 were fused to mitochondrial targeting sequences (MTS), while their C-termini were fused to EGFP to enable visualization of their cellular localization. The mitochondrial localization of chloride channels was assessed by Pearson correlation coefficients between fluorescent proteins and mitochondrial markers. **(B)** Confocal images of MTS fused ACR1 in different types of cells. The targeting in HeLa cells (first row), primary cardiomyocytes (second row), and primary neurons (third row) were demonstrated. TMRM or TagRFP-mito were used as mitochondrial markers. Scale bar, 5 μm. **(C)** Photocurrent spectra of chloride ion channels under light with different wavelengths (ACR1, n = 5 cells; ACR2, n = 3 cells; iC1C2, n = 24 cells). **(D)** mt-ACR1 exhibits minimal plasma membrane leaky targeting in primary cultured neurons. Light stimulation failed to inhibit action potential evoked by current injection. The green bar indicates light stimulation. **(E)** Fluorescence images of mt-ACR1 transfected primary cultured neurons stained with Fluo-4 and rhodamine 800 before and after light stimulation. The fluorescence of fluo-4 and rhodamine 800 were represented as pseudo-color. Scale bar, 5 μm. **(F)** Quantification of fluorescence changes of Fluo-4 and rhodamine 800 after light stimulation in primary cultured neurons expressing mt-ACR1 and control neurons. There was a significant increase in intracellular Ca^2+^ levels and a decrease in mitochondrial membrane potential (MMP) in mt-ACR1-expressing neurons compared to control neurons upon light stimulation. (mt-ACR1, n = 7 cells; control, n = 6 cells, Ca^2+^: *t-test*, *p* = 0.0041; MMP: *t-test*, *p* = 0.0004). **(G)** mt-ACR1 exhibits minimal plasma membrane leaky targeting in primary cultured cardiomyocytes. Light stimulation failed to inhibit spontaneous action potential. **(H)** Fluorescence images of mt-ACR1 transfected primary cardiomyocytes stained with Fura-2 before and after light stimulation. The calcium images were represented as pseudo-color. Scale bar, 5 μm. **(I)** Spectrogram of cardiomyocytes expressing mt-ACR1 upon light stimulation. The green bar indicated light stimulation. Continuous light stimulation can induce the cessation of beating. **(J)** Confocal images of mt-ACR1 transfected primary cardiomyocytes stained with rhodamine 800 before and after light stimulation. The images of rhodamine 800 fluorescence were represented as pseudo-color. Scale bar, 5 μm. **(K)** Quantification of rhodamine 800 fluorescence changes after light stimulation in primary cardiomyocytes expressing mt-ACR1 and control cells. Light stimulation led to a significant decrease of MMP in mt-ACR1 cells relative to control (n = 6 cells, *t-test*, *p* < 0.0001).

A sustained light illumination using ACR1 inhibited neuron firing (Figure S7C), while mtACR1 could not (Figure 3D), indicating a low level of membrane leaky expression of mt-ACR1. Interestingly, a slight shift of resting potential of neurons expressing mt-ACR1 was observed under light (Figure 3D). Light illumination also induced an elevation in calcium level and a decrease of mitochondrial membrane potential, as indicated by Fluo-4 and rhodamine 800, respectively (Figure 3E-F). Similar to neurons, mtACR1 did not inhibit cardiomyocyte beatings, but it caused a shift of resting potential (Figure 3G, Figure S7D). In mtACR1-expressed cardiomyocytes, prolonged light stimulation led to the arrest of cardiomyocytes’ beatings and calcium overload, as indicated by the calcium indicator, Fura2-AM (Figure 3I). The calcium signals were subjected to short fast Fourier transformation (SFFT)^39^ to demonstrate the frequency change upon light stimulation (Figure 3J). The result indicated that prolonged light stimulation could lead to the cessation of beating in cardiomyocytes. Additionally, light illumination caused a significant depolarization of mitochondria membrane potential as indicated by rhodamine 800 (Figure 3J-K; light vs. dark, p= 0.0009, *t-test*). Furthermore, cell death is also increased upon light stimulation as indicated by PI staining in mt-ACR1 stably transfected HeLa cells (Figure S8).

MiniSOG is a genetically encoded fluorescent protein derived from the phototropin 2 of *Arabidopsis thaliana*. When illuminated with blue light, MiniSOG efficiently generates singlet oxygen (^1^O_2_)^33^. The fluorescence of MiniSOG and its capability of ^1^O_2_ generation can be harnessed for correlated light and electron microscopy (CLEM) imaging, allowing for the localization of genetically tagged proteins in cells, tissues, and even in multicellular organisms such as intact nematodes and mice^40,41^. To explore the ROS generation effect of miniSOG on the survival of cells, miniSOG is synthesized according to the mammalian codon usage and expressed in mammalian cells. MiniSOG was targeted to mitochondria by fusing its N-terminal with a mitochondrial signal peptide (4cox8). Both cytosolic and mitochondrial miniSOG-expressing cells were stained with the mitochondrial potential indicator, TMRM, and then illuminated with blue light for 5 minutes to generate ROS (Figure 4). Interestingly, cells transfected with mitochondrial miniSOG lost their MMP upon light stimulation, while cells with cytosolic miniSOG expression maintained their MMP (Figure 4A-B; mt-miniSOG, light vs. dark, *p* < 0.0001, *t-test*; miniSOG, light vs. dark, *p* = 0.5223, *t-test*). Parkin, an E3 ubiquitin ligase, plays a pivotal role in the cellular process of mitophagy. The translocation of Parkin to mitochondria is a critical step in the initiation of mitophagy^42^. We further investigated the downstream effects of mitochondrial depolarization induced by mt-miniSOG and observed that parkin was translocated to mitochondria, which is a signal for mitophagy induction (Figure 4C). The Pearson’s correlation coefficient between the mitochondria and the fluorescence of Parkin was used to quantify the extent of parkin translocation to mitochondria, the results showed a significant portion of Parkin were translocated to mitochondria upon light stimulation (Figure 4D, pre light vs. post light, *p* = 0.0001, *t-test*).

**Figure 4.**
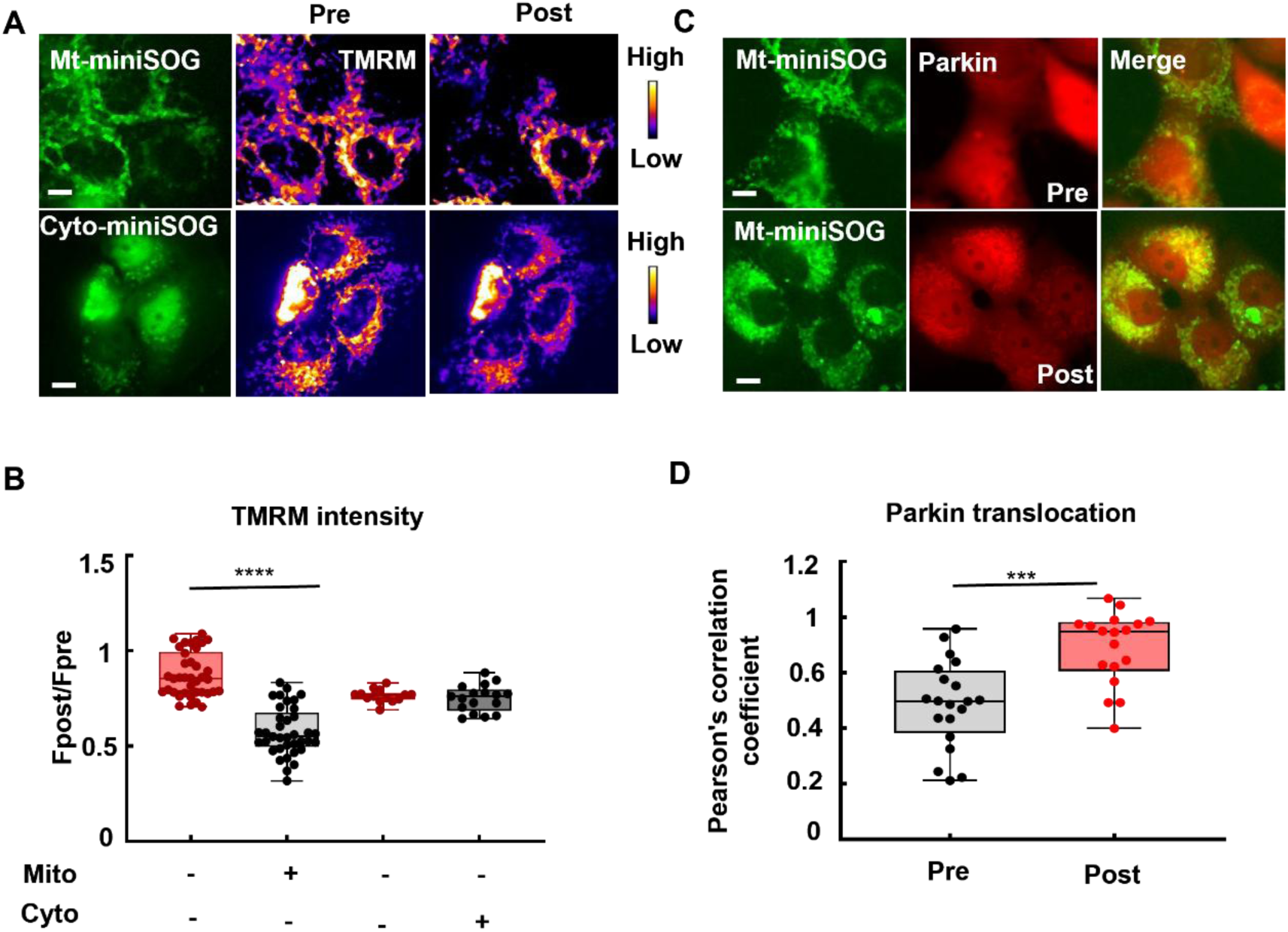
Mitochondria-targeted ROS generator induced mitochondrial depolarization and parkin translocation. (**A**) Fluorescence images of HeLa cells expressing mitochondria-targeted miniSOG and cytosolic miniSOG stained with rhodamine 800 before and after light stimulation. The fluorescence of miniSOG was represented as green, and the fluorescence of rhodamine 800 was represented as pseudo-color. The time period for light stimulation is 5 min. Scale bar, 5 μm. (**B**) Comparison of TMRM fluorescence intensity changes after stimulation in HeLa cells expressing cytosolic and mitochondrial miniSOG. Mitochondrial miniSOG caused a significant drop in MMP compared to control (mt-miniSOG, n = 37 cells; control, n = 38 cells, *t-test*, *p* < 0.0001), while cytosolic miniSOG showed no significant changes in MMP compared to control (cyto-miniSOG, n = 17 cells; control, n = 14 cells, *t-test*, *p* > 0.05). (**C**) Fluorescence images of Parkin-mCherry stably expressed HeLa cells transfected with mitochondrial miniSOG before and after light stimulation. Light stimulation in HeLa cells expressing mitochondrial miniSOG resulted in parkin translocation to mitochondria. Scale bar, 5 μm. **(D)** Pearson’s co-localization coefficient of parkin with mt-miniSOG before and after illumination in HeLa cells expressing mitochondria-targeted miniSOG. After illumination, the co-localization coefficient between parkin and mitochondria significantly increases (*t-test*, *p* = 0.0001, pre light, n = 20 cells; post light, n = 18 cells).

## Discussion

The maintenance of normal cell function is significantly influenced by intracellular pH (pHi). Intracellular acidification has been observed in response to various apoptotic stimuli, such as UV irradiation, arsenic, growth factor deprivation and somatostatin, typically resulting a drop of 0.3– 0.4 pH units^43^. A limited number of studies have described an intracellular alkalinization that is associated with apoptosis. However, accumulating studies reveals that intracellular alkalization is an initial step in the apoptotic pathway that takes place before caspase activation and DNA fragmentation^44^. Previous studies show that mild alkalization can enhance the activity of hexokinase, which is sufficient to acutely trigger cancer cell glycolysis^45^. However, increasing the cellular pH using bicarbonate reduced the pH gradient, membrane potential, and proton motive force across the inner membrane of mitochondria, which results in an increase of AMP and mitochondrial damage^46^. Archaerhodopsin (ArchT), which is a light-activated proton pump, increases the intracellular pH (pH_i_) and increases membrane dynamics, including more protrusion and retraction events^47^. In this study, GR, which acts as a light-driven proton pump, was used to study the effects of light-induced intracellular pH changes on cell metabolism and survival. Light stimulation of GR-expressing cells led to an increase in intracellular pH and a decrease in intracellular ATP levels. Our findings indicate that light can inhibit ATP generation via glycolysis and oxidative phosphorylation pathways. Moreover, light triggered cell death in cardiomyocytes expressing GR. As MPTP is opened upon alkalinization^10,46^, elevated pH in GR-expressed cells may activate MPTP and induces cell death.

Mitochondrial membrane potential (Δψ_m_) plays a crucial role in cell death. Loss of Δψ_m_ can occur either as an early event during apoptosis or as a consequence of the apoptotic signaling pathway, depending on the type of apoptotic stimuli^18^. Furthermore, decreased MMP initiates pathways to eliminate damaged mitochondria via mitophagy^48^. Previous studies demonstrated that introducing channelrhodopsin to mitochondria enables light to control MMP and influence spontaneous beats in cardiac myocytes, as well as glucose-dependent ATP increase in pancreatic β-cells^49^. Mitochondrial targeted channelrhodopsin was also used to mediate cell death in HeLa cells^50^. However, the function of chloride channels, as well as reverse proton pump on mitochondria have not been investigated. Under physiological conditions, mitochondrial chloride ion channels, such as the inner membrane targeted CLC5^38^ and the outer membrane targeted CLC4^20^, are involved in modulating mitochondrial ROS generation and induction of cell death^51^, respectively. Once chloride channels are opened, the membrane potential will shift towards the reverse potential of chloride, which depends on the concentration of Cl^-^ ions on either side of the membrane, as described by the Nernst equation. The intracellular chloride concentration varies among different cell types and can be influenced by various factors, while the extracellular chloride concentration remains around 100 to 120 mM. In hepatocytes, for instance, the intracellular chloride concentration is about 105.2 ± 62.4 mM and the chloride concentration in mitochondria was found to be 4.2 ± 3.8 mM^52^. Given that extracellular chloride is 100 mM, the chloride reverse potential on plasma membrane and inner mitochondria membrane is about 1.35 mV and -86 mV, respectively. Considering the mitochondrial membrane potential, which ranges between -180 mV and -220 mV, opening chloride channels on mitochondria can lead to the depolarization of mitochondria. Based on this calculation, it is speculated that a mitochondrial targeted light-gated chloride channel could cause mitochondrial depolarization upon light stimulation. This speculation was validated with our mt-ACR1, which depolarized mitochondria and caused calcium overload in cardiomyocytes. Consequently, mitochondria depolarization caused by mt-ACR1 might open the MPTP, resulting in calcium influx and subsequent cell death. In contrast to channelrhodopsin, which is a cation channel, mainly Na^+^, the sodium reverse potential on mitochondria is around 7.6 mV, considering the concentrations of sodium in mitochondria and cytosol are 6 mM and 8 mM, respectively^53^. While mitochondrial channelrhdopsin leads to a complete loss of MMP^54^, the mitochondrial ACR induces a mild depolarization and depends on the cell types.

The mitochondrial proton gradient is a crucial player in energy production in eukaryotic cells, where pumping of H^+^ ions across the inner mitochondrial membrane creates an electrochemical gradient to drive ATP synthesis through oxidative phosphorylation, generating vast amounts of energy for cellular activities. Depletion of this proton gradient leads to reduced ATP production, increased heat production, and ROS production. Mitochondrial uncoupling proteins, which create a channel for protons to bypass the ATP synthase and dissipate as heat, can increase energy expenditure and promote weight loss. Genetic manipulations that induce an increase in UCP1 expression can lead to a reduction in obesity and an improvement in insulin sensitivity^55^. mtASR(D217E), similar to UCPs, may provide potential treatment for obesity. In contrast to proton pumping rhodopsins, reverse proton pump acidifies the cytosol when it is expressed on plasma membrane, which also makes it a potential tool for light-induced acidification. Mild acidification is known to induce the expression of heat shock proteins^56^, protecting cells from various forms of stress. Mild acidosis also protects neurons during oxygen and glucose deprivation by attenuating the loss of mitochondrial respiration^57^. Additionally, mild acidification promotes autophagy^58^, a cellular process that eliminates damaged or dysfunctional cellular components and promotes cell survival.

Reactive oxygen species (ROS) are a group of highly reactive molecules derived from molecular oxygen capable of causing oxidative damage to biomolecules. Depending on their type, source, location, concentration, and duration, ROS can have both protective and detrimental effects on cells ^59^. When miniSOG is in mitochondria, it induces mitochondrial depolarization, while having limited effects on MMP when it is in the cytosol. This was in line with a previous study of miniSOG in *C.elegans* which demonstrated that mitochondrial outer membrane targeted miniSOG could lead to highly efficient cell killing, whereas the free form of miniSOG could not ^60^. Mitophagy selectively removes damaged mitochondria. ROS generation, induced by mt-miniSOG, could activate mitophagy through Parkin recruitment. Targeting ROS-mediated signaling pathways or modulating ROS levels may hold therapeutic potential for cancer prevention and treatment ^61^.

Overall, this paper discussed the strategies manipulating mitochondrial function for inducing cell death utilizing light-sensitive proteins. These approaches encompass several strategies, such as the inhibition of oxidative phosphorylation through GR-induced alkalization, mitochondrial depolarization via mt-ASR(D217E) and mt-ACR1, and ROS generation with mt-miniSOG. While optogenetic methods offer higher specificity than traditional drugs, it is still challenging to target only cancer cells without affecting healthy cells. Another challenge is to overcome several physiological barriers, such as tissue penetration and delivery, to reach tumor cells. Developing more efficient delivery methods, such as using nanoparticles^62^, could improve the efficacy of these methods. In conclusion, while optogenetic methods offer promising avenues for inducing cell death in cancer cells, there are still several challenges that need to be addressed. Improving the specificity, efficiency, and delivery of light-activated proteins could lead to more effective and targeted potential applications, such as cancer therapies.

## Material and Methods

### Plasmid construct

The genes GR (NCBI Accession ID: WP_011140202), BR (NCBI Accession ID: P02945), ASR (Anabaena (Nostoc) sp. PCC7120, ID: 3984), PoXeR (NCBI Accession ID: WP_051881467) and NsXeR (NCBI Accession ID: EGQ43296) and miniSOG (NCBI Accession ID: AGE44112) were synthesized according to human-codon usage. iC1C2, GtACR1 and GtACR2 were obtained from Addgene (plasmid IDs: 55630, 67795 and 67877). AT1.03 was received as a gift from Hiromi Imamura, while super ecliptic pHluorin was gifted by Gero Miesenböck. To perform light-induced pH measurements, super ecliptic pHluorin was fused with the C-terminal of GR. For light-induced ATP measurement, AT1.03 was fused with the C-terminal of GR. Furthermore, to achieve mitochondrial targeting, a four-time repeated signal sequence of cytochrome c oxidase (cox8, NP_004065, 1-29aa) was constructed using PCR and linking methods. The corresponding DNA sequence of 1-135aa mouse ABCB10 (Gene ID: 56199) was cloned from an mRNA library. Site-directed mutagenesis was conducted via a two-step megaprimer PCR method, as previously described. For expression in *E. coli*, ASR(D217E) or ASR(D217N) were cloned into the pet28c(+) vector. For current recording of ASR(D217E), super ecliptic pHluorin or AT1.03 was fused to the C-terminal of super ecliptic pHluorin to monitor expression. To express in cardiomyocytes, the cTnT promotor (-1 - -589) was amplified from the mouse genome. Furthermore, the hSyn promoter was amplified from plasmid (Addgene ID 50465) for expression in neurons. Genes were cloned into pAAV-MCS vectors for AAV virus packaging and GR-EGFP was cloned into a pCDH-CMV-puromycin vector using the restriction sites Nhe I and Not I for lentivirus packaging.

### Cell culture and transfection

HEK293t, COS7, and HeLa cells were grown in Dulbecco’s modified Eagle (DMEM, Gibco) culture medium with 10% fetal bovine serum. These cells were transfected using the calcium phosphate precipitation method and maintained at a temperature of 37°C under an atmosphere of 5% CO_2_/95% air.

### Living cell imaging

Cells were visualized using either a fluorescence laser scanning confocal microscope (FV1000, Olympus) or a fluorescence microscope (Olympus IX83, Japan). For pH imaging, cells were loaded with SNARF-1-AM (Invitrogen) to a final concentration of 20 μM in Tyrode’s solution for 20 min at 37℃. SNARF-1AM was excited by 559 nm and emission was collected at 580-600 nm and 640-700 nm. The pH was determined by calculating the ratio of the two emission channels. For ATP imaging, AT1.03 was excited by 440 nm and emission was collected at 460-490 nm and 520-550 nm. The ATP was determined by calculating the ratio of the two emission channels. Inhibition of glycolysis was achieved through treatment with 2DG (10 mM), while oligomycin (100 ng/mL) inhibited oxidative phosphorylation. To visualize cell death, PI was loaded to a final concentration of 1 μM in Tyrode’s buffer and imaged during exposure to PI solution. PI was excited by 559 nm and emission was collected at 580-630 nm. For light induced mitochondrial membrane potential imaging, cells were loaded rhodamine 800 (50 nM) or TMRM (20 mM) in Tyrode’s solution for 20 min at 37℃. Rhodamine 800 was excited by 633 nm and emission was collected at 650-750 nm. TMRM was excited by 559 nm and emission was collected at 580-630 nm. Fluo-4 (Invitrogen) was added to neurons to achieve calcium imaging, with a final concentration of 5 μM in Tyrode’s buffer for 15 minutes at 37℃. Fluo-4 was excited by 488 nm and emission was collected at 490-540 nm. For calcium imaging in cardiomyocytes, cells were loaded with Fura-2 (Invitrogen) to a final concentration of 5 μM with 0.04% F127 (Sigma-Aldrich) in Tyrode’s buffer for 15 min at 37℃. Fura2 was excited by 340 nm and 380 nm, and emission was collected at 520 ± 20 nm. The [Ca^2+^] was determined by calculating the 340 nm/380 nm ratio of the fluorescence channels.

### Light stimulation

To achieve region-specific light stimulation, a fluorescence laser scanning confocal microscopy (FV1000, Olympus) was utilized, while wide-field light stimulation was performed using a fluorescence microscope (Olympus IX83, Japan). For measuring light-induced pH changes with SNARF, a region of interest (ROI) containing a single cell was selected on the confocal image. The selected cell was then illuminated with confocal 515 nm laser. Similarly, for light-induced ATP change measurement, a ROI containing a single cell was selected on the confocal image and illuminated with confocal 559 nm laser by bleaching mode. Mitochondrial membrane potential imaging was also achieved via selection of a ROI containing a single cell on the confocal image, with subsequent illumination through confocal 515 nm laser. In contrast, miniSOG was used to induce ROS generation through wide-field light stimulation, in which cells were exposed to 488 nm light through a 30x objective for 5 minutes.

### ATP assay

HeLa cells transfected with GR-AT1.03 or GR(D121N)-AT1.03 were grown in 10% FBS DMEM until confluency reached 70%-80%. The cells were then transferred to Tyrode’s solution and exposed to green light (532nm, 37.18 mW) at room temperature (25°C) for two hours. ATP contents were determined using the ATP Assay Kit (Beyotime, China), following the manufacturer’s instructions. The experiments were performed four times and the results were presented as a box and scatter plot.

### Virus preparation and transduction

To transfect cultured cardiomyocytes, we employed the AAV DJ serotype virus. Producing this virus involved co-transfecting gene plasmids, capsid (pAAV-DJ), and helper plasmids (pHelper) into 293t cells using the calcium phosphate precipitation method. After 48 hours of incubation, viruses were collected via four cycles of frozen-thaw methods from cell pellets. Lentiviruses, on the other hand, were generated from 293t cells that were transfected with gene plasmids, capsid (pMD2.G), and helper plasmids (psPAX) via the same calcium phosphate precipitation method. These viruses were then harvested from the cell medium and concentrated by utilizing a 100 K ultrafiltration membrane (millipore, USA). Finally, the virus titer was determined through real-time PCR.

### Stable transfection cell line construction

Stable transfected cell lines were established through lentivirus transduction and subsequent screening. The virus was added to the culture medium, and cells were cultured for three days to allow for virus infection and expression. To maintain stable expression of the target gene during screening, Puromycin (20 μg/mL) was added to the medium. To confirm that plasmid sequences were present within the stable transfected cell lines, genomic DNA was extracted and subjected to PCR and sequencing analysis performed by Genewiz.

### OCR recording

An O2K system (OROBOROS) is used to measure oxidative consumption. HEK293t cells that were stable-transfected with GR-EGFP were grown in Dulbecco’s modified Eagle (DMEM, Gibco) culture medium containing 10% fetal bovine serum and harvested through centrifugation at 400g. Non-transfected HEK293t cells were used as a control. The cell pellets were then resuspended in Tyrode’s solution. Subsequently, the cells were transferred to the O2K chamber and recording began until the system was balanced. For light stimulation, a customized LED illumination device controlled by Arduino was utilized.

### Primary cardiomyocyte and neuron culture

Postnatal day 0 (P0) C57BL mice were used for either heart or hippocampi dissection. The tissues were chopped into small pieces and washed with HBSS. For heart tissues, trypsin (Sigma-Aldrich) was applied for 5 minutes followed by collagenase II (Sigma-Aldrich) for an additional 30 minutes. The cells were then dissociated further using fire-polished pipettes before being plated onto coverslips (Glasswarenfabrik Karl Hecht, Germany, 12mm) coated with matrix gel (Corning). Cardiomyocytes were cultured in plating medium and regularly fed twice a week. After 24 hours, the cardiomyocytes began to beat spontaneously. For hippocampal tissues, trypsin was used to digest the cells for five minutes, after which single cells were produced through further dissociation via fire-polished pipettes. The cells were then harvested by centrifugation and plated onto matrix gel coated coverslips. They were also cultured in plating medium and fed twice a week.

### Electrophysiology

A customized opto-electro system was used to perform whole-cell patch clamp recordings at room temperature. The system consisted of an amplifier (Axopatch 700B, Molecular Devices), a digitizer (Digidata 1440A, Molecular Devices), a signal generator (Master 9), a laser system with 470 nm, 532 nm and 633 nm lasers, a monochromator (Optoscan, Cairn Research Ltd., UK) with a Xenon bulb, and an imaging system (Olympus). Micro-Manager was used for device control, and data were acquired at a sampling rate of 10 kHz. Micropipettes with a tip resistance of 4-7 MΩ were pulled from filamented glass capillaries (Sutter Instrument, BF150-86-10) using a micropipette puller (Sutter Instrument, P1000) filled with intracellular buffer (potassium gluconate 120 mM, KCl 3 mM, HEPES 10 mM, NaCl 8 mM, CaCl_2_ 0.5 mM EGTA 5 mM, ATP-Mg 2 mM, GTP 0.3 mM, pH 7.2) and positioned using a micromanipulator (Sutter, MP285). For light-activated photocurrent recording, HEK 293t cells were recorded at voltage clamp mode, and photocurrents were evoked by 532 nm laser illumination through a 40x objective (Olympus). Action spectrum recording was performed by triggering the monochromator to generate 300 ms pulses of light with a 20 nm bandwidth and wavelength stepping from 350 nm to 750 nm, with light originating from the Xenon bulb. For recording I-V curve, a protocol was set to hold the cells at a membrane potential from 40 mV to -80 mV with a -20 mV voltage step, accompanied by a digital signal to trigger the 532 nm laser for generating green light pulses (130 mW mm^-2^, interval: 1 s, duration: 1s). Action potential recording was performed in primary neurons and cardiomyocytes. Cultured neurons were transfected by calcium phosphate precipitation method at days in vitro (DIV) 4-5 and recorded at DIV 8-12. Neurons were voltage clamped at -70 mV and recorded at current clamp mode. For action potential recording in primary cardiomyocytes, cultured cardiomyocytes were transfected by AAV virus at DIV 2-3 and recorded at DIV 7-10. Light pulses and current pulses were generated using customized protocols.

### Colocalization analysis

Fluorescence laser scanning confocal microscopy (FV1000, Olympus or LSM980, Zeiss) was used to capture images. To assess mitochondrial colocalization, we cropped the confocal images in Image J (NIH) to a small area containing only one cell. Colocalization analysis was performed using the Plugin Colocalization Indices, and the effectiveness of mitochondrial targeting was scored using Pearson’s correlation coefficients.

### Heterologous expression in *E.coli*

ASR(D217E) and ASR(D217N) were cloned into pet28c(+) vectors for expression. Clones of *Escherichia coli* (strain BL21, DE3) transformed with expression plasmids are inoculated and grown in 2 ml (small scale) or 100 ml (large scale) of Luria-Bertani (LB) medium supplemented with 30 μg/ml kanamycin at 180 rpm, 37 °C. Growth was monitored using optical density measurements at 600 nm until the cells reached an OD600 of 0.8. Subsequently, 10 μM of all-trans retinal and 1 mM of IPTG were added, and the cells were allowed to grow for an additional 18 h at 28°C while being shaken at 180 rpm. The cells were eventually harvested through centrifugation, and their color changes were observed to confirm successful expression of the microbial rhodopsins.

### pH recording in *E.coli* suspension

*E. coli* cells expressing ASR(D217E) and ASR(D217N) were harvested by centrifugation at 8000 g for 10 min at room temperature, and then resuspended in a solution containing 150 mM NaCl and 50 mM MgSO_4_, to a concentration of 0.1 g/ml. The *E. coli* suspension was subjected to illumination with a 532 nm laser, while pH changes were monitored using a Micro-Combination pH Needle Electrode (Microelectodes) connected to an A/D converter, Digidata 1440 A. Temperature signals were recorded using a temperature sensor that was connected to a temperature controller (Warner). Data acquisition was carried out using AxoScope 10.4 software (Axon Instruments) at a sampling rate of 1000 Hz. The pH signals were calibrated for temperature using the following equation:

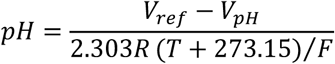

where V_ref_ is the standard electrode potential when pH is 7.0, V_pH_ is the voltage measured with the pH electrode, R is the gas constant, T is the recorded temperature in Celsius, and F is the Faraday constant.

The *E. coli* suspension was exposed to three green pulses (532 nm, 37.18 mW cm^-2^, pulse duration: 2 min, pulse interval: 2 min) during recording.

### Video-based cardiomyocytes beating analysis

Based on the Nyquist sampling theorem, the sampling frequency for imaging must be no less than twice the beating frequency of cardiomyocytes. In this experiment, bright field images were captured via an EMCCD camera (Photometric Evolve) at a sampling rate of 8 Hz, then analyzed using ImageJ (NIH). Firstly, the video was converted to 8-bit, then a line covering cell movement was drawn on it to generate a kymograph by Multiple Kymograph plugin. A plot of grayscale-value versus time was obtained by drawing a straight line on the time-axis of the kymograph with the “Plot Profile” command. The grayscale-value data was saved and used for spectrogram analysis.

### Parkin translocation assay

Parkin-mcherry stable-transfected cells were subjected to lentivirus transduction for mt-miniSOG expression. Following this, the cells were illuminated with light for 30 min and cultured in dark for additional 1 h. Then, cells were imaged to assess puncta accumulation. The colocalization of mt-miniSOG and parkin was analyzed before and after light illumination.

### Data and statistical analysis

All electrophysiology data were analyzed using pClampfit (Molecular Devices) in combination with customized Perl scripts. Statistical analysis was performed using unpaired two-tailed Student’s t tests being used for comparison of two samples.

## Author Contribution

Conceptualization: R.Z.Y, J.S.K; Data curation: R.Z.Y; Formal analysis: R.Z.Y; Funding acquisition: P.P.L, S.A.L, J.S.K; Investigation: R.Z.Y; Methodology: R.Z.Y, D.D.W, S.M.L, D.H.L; Project administration: J.S.K; Resources: P.P.L, S.A.L; Software: R.Z.Y; Supervision: J.S.K; Validation: R.Z.Y; Visualization: R.Z.Y; Writing – original draft: R.Z.Y; Writing – review & editing: R.Z.Y, J.S.K.

