## supplementary file for "Optogenetic Regulation of Mitochondrial Function to Modulate Cell Death"

**Supplementary Information**

**Supplementary Figure 1. Proton pump characteristics of GR.** (**A**) Fluorescence changes of GR-pHluorin expressing HeLa cells upon light stimulation. The red curve represents GR-pHluorin expressing HeLa cells with light stimulation, while black curve represents the cells without light stimulation (light, n = 3 cells; control, n = 3 cells). The light stimulation condition was 515 nm, 60 s. (**B**) pH curve of *E.coli* suspension expressing GR upon light stimulation. The green bars indicate light stimulation.

**Supplementary Figure 2. Light-induced intracellular ATP reduction by BR.** (**A**) Patch clamp recording of photocurrent of HEK293t cells expressing Bacteriorhodopsin (BR). The green bar indicates light stimulation. (**B**) Fluorescence images of BR expressing HeLa cells stained with SNARF before and after light stimulation. The images were presented as SNARF ratios in pseudo-color. The white dashed circle in the figure indicated the region with light stimulation. The light stimulation condition was 515 nm, 60 s. Scale bar, 5 μm. (**C**) Curves of SNARF ratios in BR-expressing HeLa cells and control cells upon light stimulation. Red curve represents HeLa cells transfected with BR, while black curve represents the control cells (BR, n = 4 cells; control, n = 4 cells). The light stimulation condition was 515 nm, 60 s. (**D**) Fluorescence images of HeLa cells expressing the fusion protein BR-AT01.03 before and after light stimulation. The images were presented as AT1.03 ratios in pseudo-color. The white dashed circle in the figure indicated that the region with light stimulation. The light stimulation condition was 559 nm, 60 s. Scale bar, 5 μm. (**E**) Curves of AT1.03 ratios in BR-expressing HeLa cells and control cells upon light illumination. Red curve represents Hela cells transfected with BR-AT1.03, while black curve represented the control cells (BR, n = 4 cells; control, n = 4 cells). The light stimulation condition was 559 nm, 60 s.

**Supplementary Figure 3. Intracellular targeting of mt-NsXeR-EGFP in HeLa cells**. Confocal images of mt-NsXeR-EGFP in HeLa cells. TMRM was used as a mitochondrial marker. mt-NsXeR-EGFP in HeLa cells demonstrated non-mitochondrial targeting (top) and partial mitochondrial targeting (bottom). Scale bar, 5 μm.

**Supplementary Figure 4. Mitochondrial targeting of ASR(D217E).** Confocal images of mtASR(D217E)-EGFP targeting in different cells. mtASR(D217E)-EGFP localized to mitochondria in both COS7 (top) and HepG2 (bottom) cells. TMRM was used as a mitochondrial marker. Scale bar, 5 μm.

**Supplementary Figure 5. Photocurrent of PoXeR in HEK293t cells.** HEK293t cells expressing PoXeR were recorded for photocurrent using patch clamp with light stimulation. The green bar indicates light stimulation.

**Supplementary Figure 6. Photo-electrical characteristics of ASR(D217E).** (**A**) Photocurrents of ASR(D217E) at different light intensities. The photocurrents were recorded by patch clamp in ASR(D217E)-expressing HEK293t cells with light intensities ranging from 0-130 mW/cm$^2$. (**B**)

Light stimulation induced action potentials in ASR(D217E)-expressing primary cultured neurons. The red curve indicates the injected current and the green bars indicate light stimulation.

**Supplementary Figure 7. Localization of non-mitochondrial targeting chloride channels and their photo-electrical characteristics.** (**A**) Confocal images of mt-ACR2-EGFP and mt-iC1C2-EGFP in HeLa cells. TMRM was used as a mitochondrial marker. Scale bar, 5 μm. (**B**) Current-voltage (I-V) relationship of chloride channels. The red curve represents ACR1, the green curve represents ACR2, and the gray curve represents iC1C2 (ACR1, n = 3 cells; ACR2, n = 3 cells; iC1C2, n = 3 cells). (**C**) Light induced inhibition of action potential in ACR1-expressing primary cultured neurons. Action potentials were evoked with current injection (red curve). (**D**) Light induced inhibition of spontaneous action potential in ACR1-expressing primary cultured.

**Supplementary Figure 8. Mitochondrial-localized ACR1 induced cell death upon illumination**. HeLa cells stably transfected with mt-ACR1 underwent cell death as evidenced by increased PI staining after illumination (532 nm, 16 h). Scale bar, 50 μm.

Figure S1

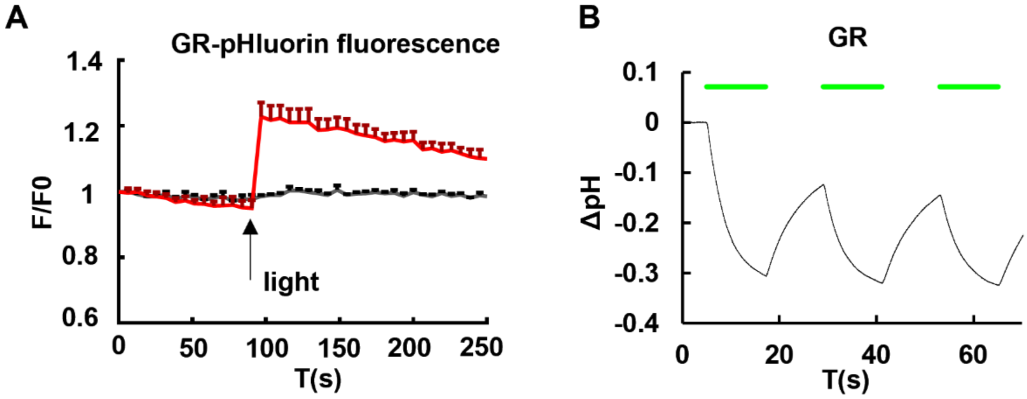
A
GR-pHluorin fluorescence
1.4
1.2
1
0.8
0.6
F/F0
light
0
50
100
150
200
250
T(s)
B
GR
0.1
0
-0.1
-0.2
-0.3
-0.4
ΔpH
0
20
40
60
T(s)

Figure S2

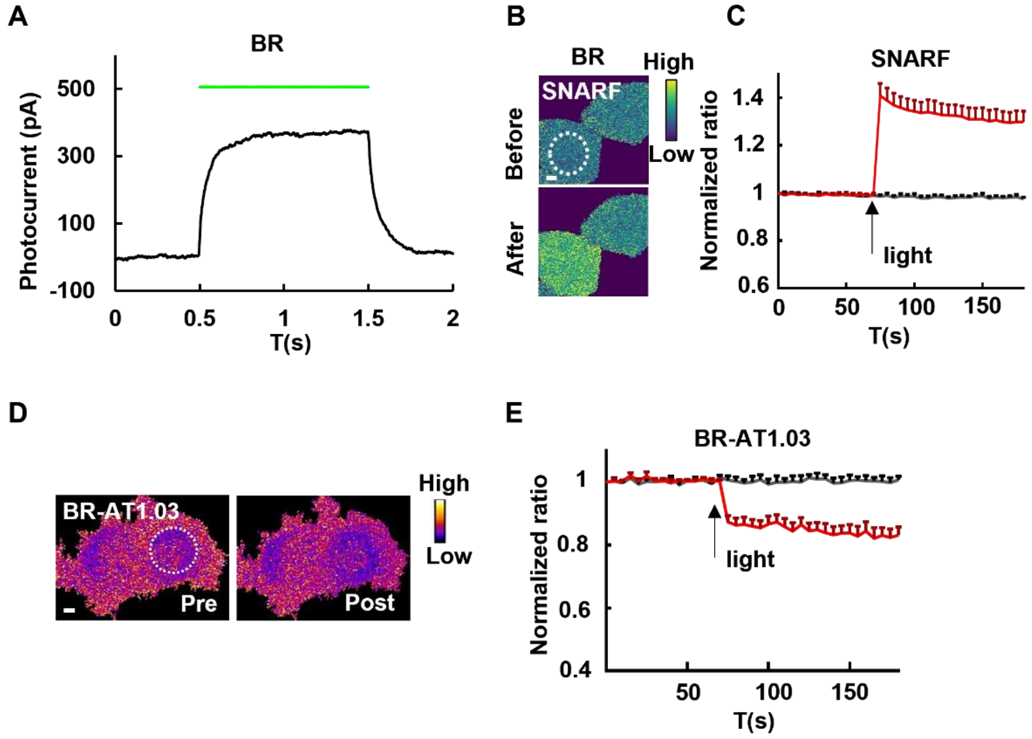
A
BR
Photocurrent (pA)
500
300
100
-100
0
0.5
1
1.5
2
T(s)
B
BR
High
Low
SNARF
Before
After
C
SNARF
Normalized ratio
1.4
1.2
1
0.8
0.6
light
0
50
100
150
T(s)
D
BR-AT1.03
Pre
Post
High
Low
E
BR-AT1.03
Normalized ratio
1
0.8
0.6
0.4
light
50
100
150
T(s)

**Figure S3**

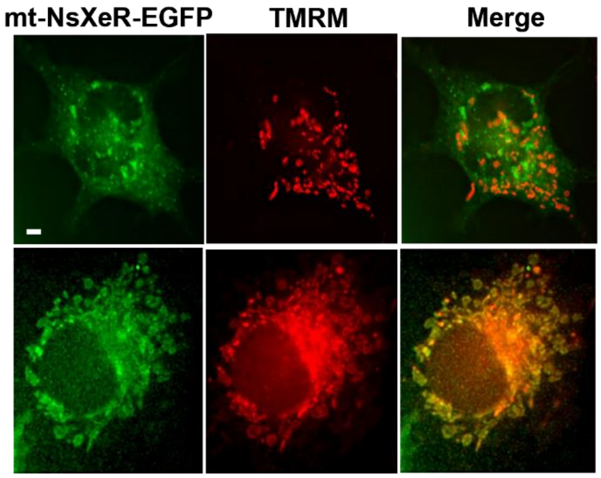

**Figure S4**

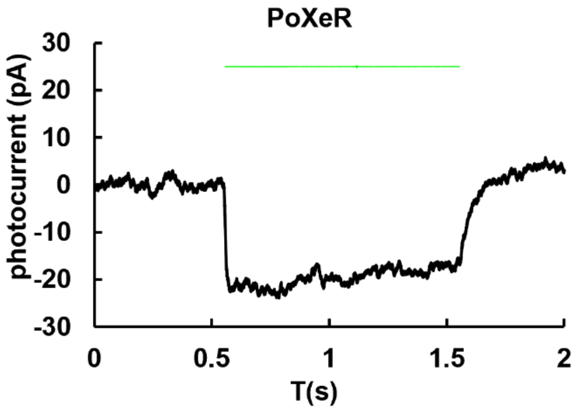

**Figure S5**

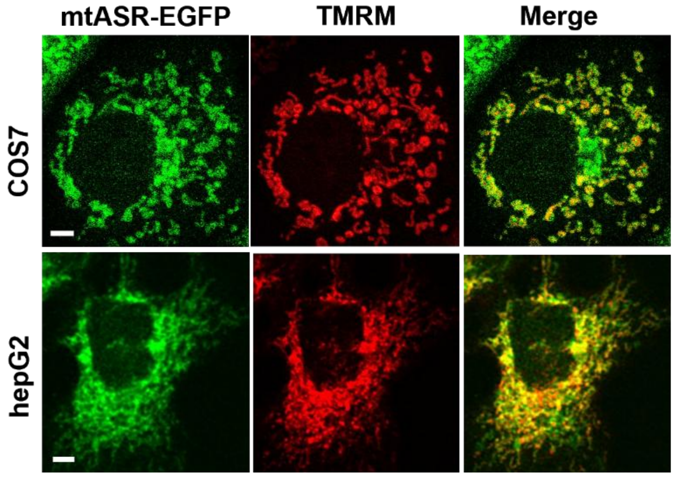

**Figure S6**

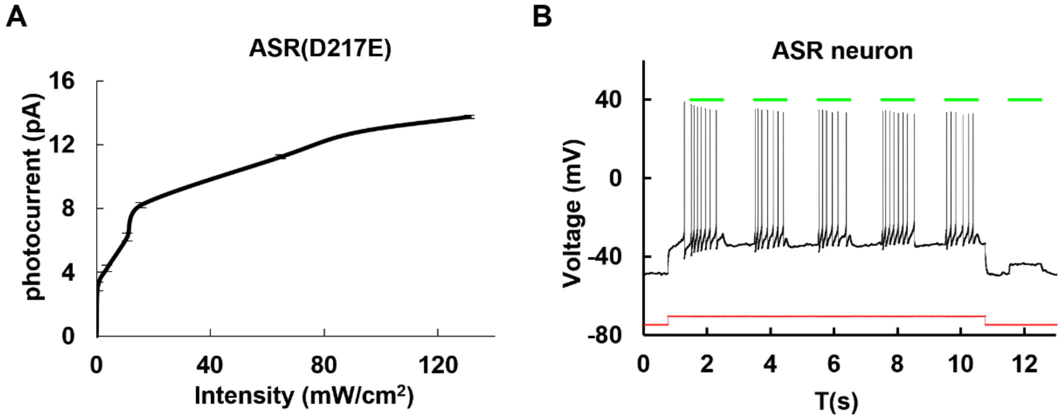

A
ASR(D217E)
photocurrent (pA)
16
12
8
4
0
0
40
80
120
Intensity (mW/cm2)
B
ASR neuron
Voltage (mV)
40
0
-40
-80
0
2
4
6
8
10
12
T(s)

**Figure S7**

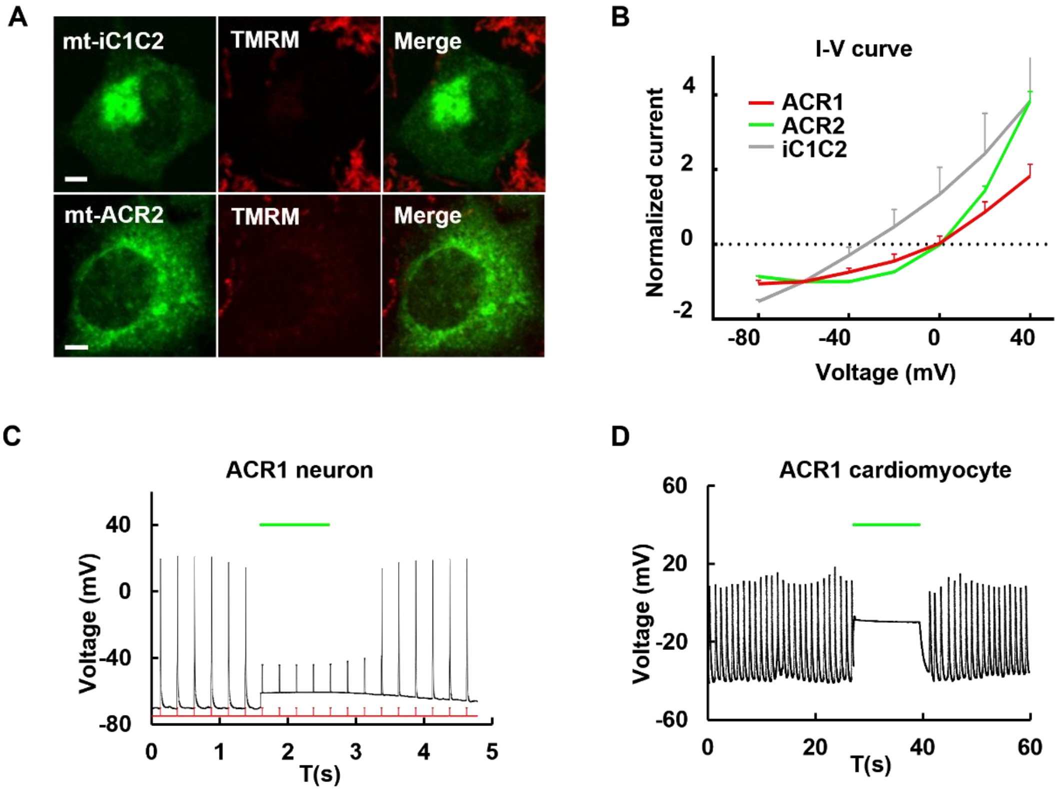
A
mt-iC1C2
TMRM
Merge
mt-ACR2
TMRM
Merge
B
I-V curve
ACR1
ACR2
iC1C2
Normalized current
4
2
0
-2
-80
-40
0
40
Voltage (mV)
C
ACR1 neuron
Voltage (mV)
40
0
-40
-80
0
1
2
3
4
5
T(s)
D
ACR1 cardiomyocyte
Voltage (mV)
60
20
-20
-60
0
20
40
60
T(s)

**Figure S8**

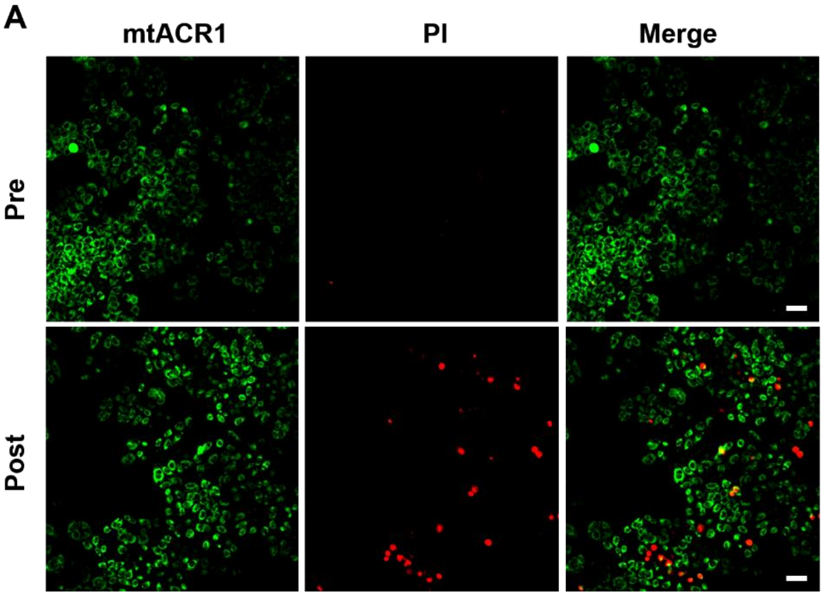
